# Subcortical volume and white matter integrity abnormalities in major depressive disorder: findings from UK Biobank imaging data

**DOI:** 10.1101/070912

**Authors:** Xueyi Shen, Lianne M. Reus, Simon R. Cox, Mark J. Adams, David C. Liewald, Mark E. Bastin, Daniel J. Smith, Ian J. Deary, Heather C. Whalley, Andrew M. McIntosh

## Abstract

Previous reports of altered grey and white matter structure in Major Depressive Disorder (MDD) have been inconsistent. Recent meta-analyses have, however, reported reduced hippocampal grey matter volume in MDD and reduced white matter integrity in several brain regions. The use of different diagnostic criteria, scanners and imaging sequences may, however, obscure further anatomical differences. In this study, we tested for differences in subcortical grey matter volume (n=1157) and white matter integrity (n=1089) between depressed individuals and controls in the subset of 8590 UK Biobank Imaging study participants who had undergone depression assessments. Whilst we found no significant differences in subcortical volumes, significant reductions were found in depressed individuals versus controls in global white matter integrity, as measured by fractional anisotropy (FA) (β=-0.182, p=0.005). We also found reductions in FA in association/commissural fibres (β=-0.184, p_corrected_=0.010) and thalamic radiations (β=-0.159, p_corrected_=0.020). Tract-specific FA reductions were also found in the left superior longitudinal fasciculus (β=-0.194, p_corrected_=0.025), superior thalamic radiation (β=-0.224, p_corrected_=0.009) and forceps major (β=-0.193, p_corrected_=0.025) in depression (all betas standardised). Our findings provide further evidence for disrupted white matter integrity in MDD.

## Introduction

Major Depressive Disorder (MDD) is a common psychiatric illness, affecting between 5 and 30% of the population which accounts for around 10% of all days lived with disability^1^. There is therefore an urgent need to identify the mechanisms underlying MDD and human *in vivo* MRI has been widely applied in this search^2^.

Many brain imaging studies have measured grey matter volume differences between healthy individuals and, predominantly clinically ascertained, individuals with MDD. Prefrontal cortex and limbic areas are fundamental to emotion processing and mood regulation^3^, and these areas have also been consistently implicated in imaging studies of MDD ^4−6^. As the use of automated methods such as voxel-based morphometry^7,8^ and Freesurfer^9^ have increased, this has expanded the search across the whole brain. In general, structural abnormalities have been reported across diverse brain networks in MDD. Regions including the thalamus^10^, amygdala^4^, insula^8^, caudate^9^, anterior cingulate cortex^4^, along with prefrontal areas such as orbital prefrontal cortex (OFC)^11^ and dorsal lateral prefrontal cortex (PFC)^12^ have been reported to be smaller in MDD versus healthy controls. However, other studies have found conflicting results^9,13^, or have reported null findings^7^. This inconsistency may be due to limited sample sizes and other sources of heterogeneity such as sample characteristics, recruitment criteria, data acquisition and image processing^14^.

The lack of a single anatomically circumscribed abnormality in MDD has led many to suggest that the disorder might be due to abnormalities of brain networks affecting connections between several regions. In support of this, findings from individual studies of white matter structure in MDD have shown patterns of alteration using diffusion tensor imaging (DTI). Proxy measures of white matter integrity, including fractional anisotropy (FA) and mean diffusivity (MD), have been used to infer connectivity differences between groups. Decreased FA indicates lower directionality of water molecule diffusion along fibre pathways and is a proxy of decreased tract integrity, whilst increased MD indicates less constrained water molecule diffusion and a proxy for lower integrity.

White matter integrity of frontal-limbic tracts have been suggested to underlie clinical features in MDD due to a lack of frontal cortical control over brain regions that involve in emotion processing^15^. Studies have reported altered water diffusivity of white matter tracts in MDD compared to healthy controls, but the tracts identified are often inconsistent. Some studies reported decreased white matter integrity in tracts that connect prefrontal areas (e.g. fronto-occipital fasciculus, superior longitudinal fasciculus)^16^. While some studies using similar sample sizes also found consistent results^17^, other groups reported FA deficits in limbic areas (e.g. posterior thalamic radiation, posterior corona radiata)^15^. Similar to the studies of subcortical volumes described above, DTI investigations of MDD have often used relatively small sample sizes^17,18^.

Meta-analytic methods may help to overcome issues related to small sample sizes and are also able to quantify and test for between-study heterogeneity. A recent meta-analysis of subcortical structures by Schmaal *et al.* tested over 1650 MDD patients and around 7000 healthy controls across 15 studies, and reported hippocampal grey matter volume reductions in MDD. No other case-control differences were found^19^. Meta-analyses of white matter integrity measures in MDD have also reported FA reductions in superior longitudinal fasciculus, fronto-occipital fasciculus, and thalamic radiations^17,18^. These studies, however, often require the combination of imaging data from different scanners, using different ascertainment criteria and methodology, different clinical instruments and have differing levels of phenotypic data to pursue further research questions. Meta-analytic findings therefore highlight the pressing need to measure brain structural abnormalities in MDD using larger single-scanner samples where robust conclusions can be made in the absence of differing study methodologies.

In the current study, we examined the volumetric structural imaging data of subcortical brain structures and tract-specific white matter integrity measures from the UK Biobank imaging study. UK Biobank is a study of 500,000 subjects recruited from across the United Kingdom^20^. The dataset used in the current study is the latest release of imaging data on 8590 participants who participated in the brain imaging assessment^21^. For our current purposes this included 354/342 MDD and 803/762 controls respectively who provided usable for T1-weighted/DTI data from a single scanner, along with available data regarding diagnostic and phenotypic information. The scanning protocol and pre-processing pipelines were devised by UK Biobank, with consistent, compatible setting of scanner parameters and participant-friendly experimental procedures. This data therefore allowed us to explore structural changes associated with depression in a single large population-based sample using data from an individual study source with unified depression classification, and with scanning sequences and image processing procedures applied consistently across all subjects, all of whom were imaged on a single MRI scanner.

## Method

### Participants

In the latest release of imaging data from UK Biobank, 5797 people completed the subcortical brain structural MRI measurements and 5171 completed DTI assessment (Fig S1). The study has been approved by the National Health Service (NHS) Research Ethics Service (approval letter dated 17th June 2011, reference: 11/NW/0382), and by the UKB Access Committee (Project #4844). Written informed-consent was obtained from each subject. All assessments were performed in accordance with the regulations and protocols from the committees.

Individuals from the initial pilot phase of imaging using different acquisition parameters were excluded from the current study, as were those that did not complete pre-processing quality checks conducted by UK Biobank. In addition, scans from individuals that were identified by our internal quality check as having a structural measure that lay more than three standard deviations from the sample mean were excluded (Fig S2, S3, Table S1). Any participants that had a diagnosis of Parkinson’s Disease, bipolar disorder, multiple personality disorder, schizophrenia, autism or intellectual disability were also excluded from the current analysis (ICD-10/9 or self-report). This resulted in data from 5397 participants with T1-weighted subcortical volumes and 4590 participants with DTI measures. Mean ages were 55.47 +/- 7.49 years for those with T1-weighted, grey matter data and 55.46 +/- 7.41 years for those with DTI, white matter integrity. The proportions of male participants are were similar in both datasets (45.78% for those providing T1-weighted data and 47.12% for those with DTI measures). Details of data exclusions are detailed within supplementary materials (Method, Participants; Fig S1).

### MDD definitions

The definition of MDD used in the current study was generated based on the putative MDD category summarized previously by Smith *et al.*, as presented in supplementary materials (Fig S4)^22^. They generated the criteria of single episode major depression, recurrent major depression (moderate), recurrent major depression (severe) and those who were absent of depression. This category was benchmarked by testing its prevalence in the sample, and by testing for association with a number of traits, such as neuroticism^23^, that have previously been associated with MDD^24^. However, since the category is based on hospital admission data and depressive symptoms, which were both self-reported, rather than more formal ICD/SCID criteria, cases should be considered ‘probable’ MDD rather than operationally defined on the basis of an interview.

We generated two definitions of probable MDD. One was the principal MDD definition that compared all MDD patients (recurrent and single episode) with healthy controls, while the other was based on recurrence and compared recurrent MDD patients with non-recurrent and non-MDD individuals.

The principal MDD definition therefore included those who were categorised in single and multiple episode major depression as cases. The corresponding control group contained participants that were absent of depression according to the putative MDD category described by Smith *et al*^22^. For the recurrent MDD definition, the case group only included recurrent major depression. The corresponding control group therefore referred to the participants without recurrent MDD, which included single episode major depression, those who were absent of depression and those who reported depressive symptoms but not enough to be specified as MDD. Participants who did not answer one or more of the questions necessary for classification were excluded from this analysis.

For each definition of probable MDD, the participants with subcortical volume data consisted of 354 MDD cases and 803 controls and 261 MDD cases and 1196 controls respectively for principal and recurrent definitions. Participants with DTI data consisted of 335 MDD cases and 754 controls and 242 MDD cases and 1113 controls for principal and recurrent definitions respectively. Method used to derive the samples into analyses were presented in supplementary materials, Fig S1.

The descriptions and demographic characteristics of each MDD definition are shown in supplementary materials (Table S2, S3). For the purposes of the current analysis, we used the principal definition of depression as the main definition as it most closely resembles the general application of typical clinical criteria. We also report results of the recurrent definition of MDD to highlight differences associated with a more severe recurrent MDD diagnosis. (Supplementary materials, Table S3).

### MRI acquisition and analyses

We used the imaging-derived phenotypes (IDPs) generated by UK Biobank. The MRI acquisition, pre-processing and imaging analysis for subcortical volumes and FA values of white matter tracts were all conducted by UK Biobank using standard protocols^21^, see supplementary material. Briefly, all imaging data was collected on a Siemens Skyra 3T scanner (https://www.healthcare.siemens.com/magnetic-resonance-imaging) and was preprocessed using FSL packages. For T1-weighted data, segmentation of brain was conducted in two steps: firstly, a tissue-type segmentation using FAST (FMRIB’s Automated Segmentation Tool)^25^ was applied to extract cerebrospinal fluid, grey matter and white matter; then subcortical structures are extracted using FIRST (FMRIB’s Integrated Registration and Segmentation Tool)^26^. For DTI data, parcellation of tracts were conducted using AutoPtx^27^.

The summary data contained volumes of grey matter, white matter, cerebrospinal fluid, thalamus, putamen, pallidum, hippocampus, caudate, brain stem, amygdala and accumbens (Fig S2). DTI data provided tract-averaged FA for 27 major tracts (12 bilateral tracts in both hemispheres and 3 tracts that pass across brain): (a) association and commissural fibres: forceps major and minor, inferior fronto-occipital fasciculus, uncinate fasciculus, cingulum bundle and superior longitudinal fasciculus; (b) thalamic radiations: anterior, superior and posterior thalamic radiations; (c) projection fibres: corticospinal tract, acoustic radiation, medial lemniscus, middle cerebellar peduncle.

Scans with severe and obvious normalization problems were excluded by UK Biobank. In addition we also excluded observations that were more than three standard deviation from the sample mean for the analysis of subcortical volumes. For DTI measures, participants with at least one outlier of tract-averaged FA from the sample mean were excluded for that measure. Descriptions of the sample were reported in supplementary materials (Method, MRI preprocessing; Fig S1-3). For transparency, the results without excluding outliers are also presented in the supplementary materials.

### Statistical methods

#### Subcortical volumes

First, differences in global intracranial volume (ICV) associated with a probable MDD diagnosis were examined by modelling ICV as dependent variable, controlling for age, age^2^, sex and assessment centre. ICV was measured by adding up volumes of white matter (WM), grey matter (GM) and cerebrospinal fluid (CSF). For bilateral subcortical volumes, age, age^2^, sex, hemisphere, assessment centre and ICV were set as covariates in a repeated-effect linear model to test for an association between both probable MDD definitions on subcortical volumes, adjusted for whole brain size. For unilateral structures, a general linear model was applied as above, without controlling for hemisphere. We also examined the interaction of hemisphere and MDD definitions on bilateral structures. Where there was a significant MDD by hemisphere interaction, analyses on both lateralised structures were conducted separately. All subcortical volumes were rescaled into zero mean and unitary standard deviation in order that effect sizes represent standardized scores. False Discovery Rate (FDR) multiple comparison correction was applied for tests of the 8 subcortical volumes plus additional tests on ICV, conducted separately for the two probable MDD definitions (Fig 1, Table 1, S5).

**Figure 1.**
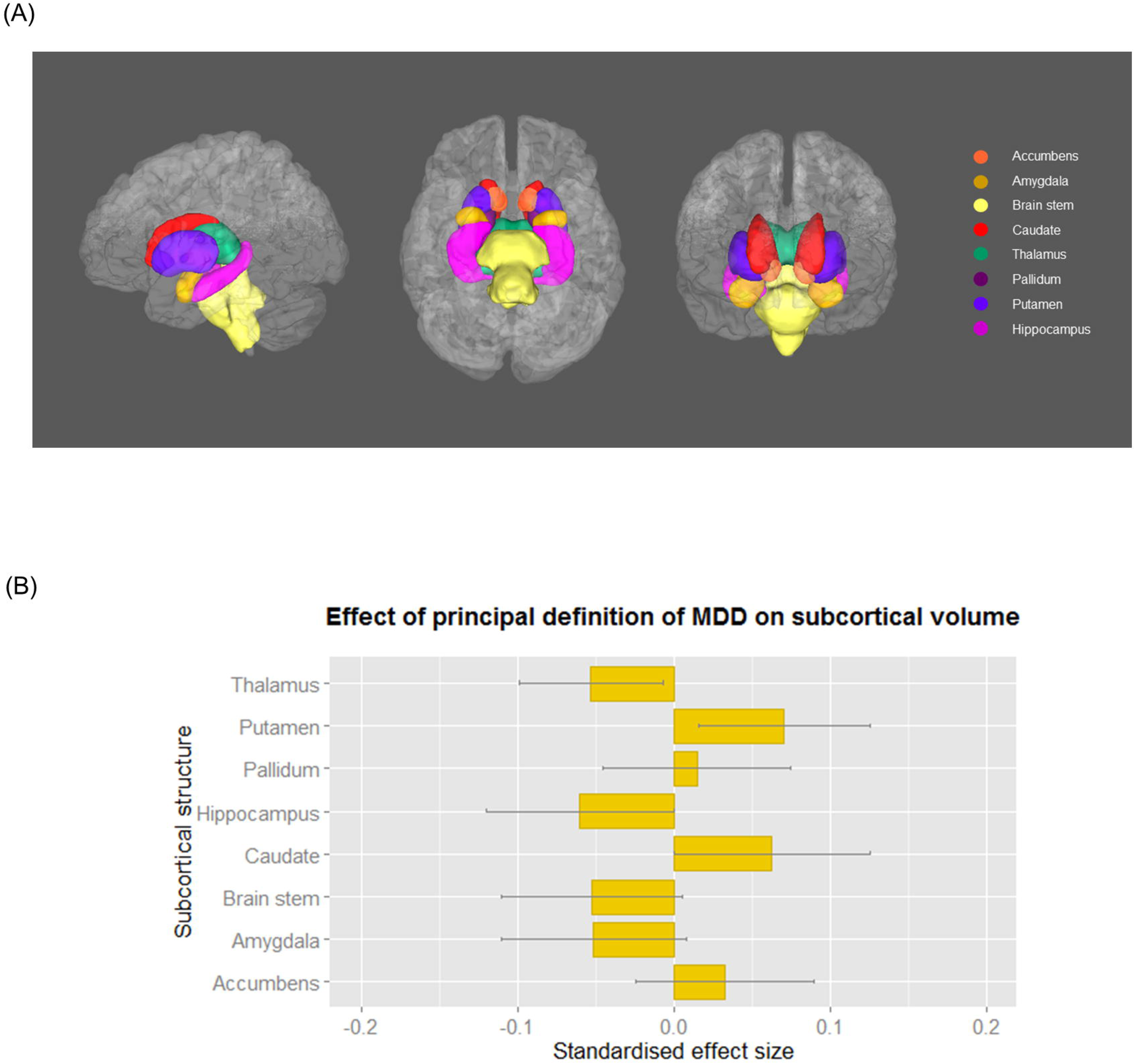
(A) Subcortical structures of interest in left, inferior and anterior view. (B) The effect of principal definition of probable MDD on subcortical volumes. Linear models were conducted, controlling the effect of age, age^2^, sex, assessment centre and intracranial volume (and hemisphere for the regions that have bilateral values). The x-axis shows the standardised effect size of MDD definition, and y-axis is the layout of the subcortical structures. The error bar represents standard deviation of mean.

**Table 1.** The effect of MDD definition on the volumes of subcortical regions and brain matters.

| Subcortical regions | Principal definition |  |  |  |  | Recurrent definition |  |  |  |  |
| --- | --- | --- | --- | --- | --- | --- | --- | --- | --- | --- |
|  | Effect size | Standard deviation | t value | p value | p<sub>corrected</sub> | Effect size | Standard deviation | t value | p value | p<sub>corrected</sub> |
| **Accumbens** | -0.010 | 0.049 | -0.211 | 0.833 | 0.838 | -0.018 | 0.052 | -0.348 | 0.728 | 0.819 |
| **Amygdala** | -0.045 | 0.050 | -0.896 | 0.371 | 0.834 | 0.038 | 0.053 | 0.711 | 0.477 | 0.819 |
| **Caudate** | 0.064 | 0.053 | 1.198 | 0.231 | 0.834 | 0.025 | 0.056 | 0.453 | 0.650 | 0.819 |
| **Hippocampus** | -0.034 | 0.050 | -0.682 | 0.495 | 0.838 | -0.040 | 0.053 | -0.758 | 0.449 | 0.819 |
| **Pallidum** | 0.019 | 0.051 | 0.372 | 0.710 | 0.838 | -0.022 | 0.054 | -0.414 | 0.679 | 0.819 |
| **Putamen** | 0.018 | 0.047 | 0.386 | 0.700 | 0.838 | -0.008 | 0.049 | -0.162 | 0.871 | 0.871 |
| **Thalamus** | -0.050 | 0.039 | -1.284 | 0.199 | 0.834 | -0.059 | 0.041 | -1.428 | 0.154 | 0.819 |
| **Brain stem** | -0.011 | 0.053 | -0.205 | 0.838 | 0.838 | 0.045 | 0.056 | 0.794 | 0.428 | 0.819 |
| **ICV** | -0.046 | 0.048 | -0.953 | 0.341 | 0.834 | -0.049 | 0.051 | -0.959 | 0.338 | 0.819 |

#### White matter integrity

In order to test for an association between probable MDD and FA, as above we used a general linear model with age, age^2^, sex and assessment centre as covariates and the definition of MDD as a fixed factor. First we examined for the effects of diagnosis on global whole brain white matter integrity. The brain’s white matter tracts have been shown to share a considerable proportion of variance in their microstructural properties in this^28^ and other samples^29,30^. Global integrity was determined using standardised approaches by applying principal component analysis (PCA) on the 27 tracts to extract a latent measure^31^. Scores of the first un-rotated component of FA were extracted and set as the dependent variable of the general linear model to test the effect of probable MDD diagnosis (variance explained=36.5%). Then we separately examined three subsets of white matter tracts: (a) association and commissural fibres which include tracts connecting cortex to cortex, (b) projection fibres which consist of tracts connecting cortex to spinal cord and brainstem, as well as sensory tracts that connect cortex to thalamus and (c) thalamic radiations that connect thalamus with cortical areas^28^. Scores of the principal un-rotated component for each subset was extracted (variance explained=44.1 %, 60.1% and 38.1% respectively for A/CF, TR and PF) for further general linear modelling as with the global latent measure. Loadings and scree plot of PCA analyses are in supplementary materials (Table S10, Fig S5). Finally, we examined the effects of depression on each tract individually. Repeated-effect linear models were used for the measures of bilateral white matter tracts correcting for hemisphere as above, while random-effect general linear models were used for the unilateral midline tracts. Both the main effect of MDD definition and its interaction with hemisphere were tested. Where the interaction was significant, tests were applied individually for left and right sides separately. FDR correction was individually applied over the three subsets of white matter tracts as well as individual tracts^32^.

## Results

### The effect of MDD definitions on subcortical volumes

We found no significant group effect for ICV based on the principal definition of MDD (β=-0.046, p_uncorrected_=0.341). There were also no significant differences between groups based on the principal definition of MDD for any of the subcortical brain regions, including the hippocampus (βs=-0.050~0.064, ps_uncorrected_>0.199, ps_corrected_>0.834); see Fig. 1, Table 1. No region demonstrated significant interaction of hemisphere, therefore no region was examined separately on different hemispheres.

The same models were also applied to compare recurrent MDD and controls, see above. No subcortical regions reached significance in this definition of recurrent cases versus controls. The largest nonsignificant effect size was observed for the caudate (β=0.064, p_uncorrected_=0.231).

### The effect of probable MDD on measures of white matter integrity

Firstly we tested the effect of probable MDD on general white matter FA (gFA). For both the principal and recurrent definitions, gFA was lower in probable MDD cases versus controls (β=-0.182, p=0.005; β=-0.160, p=0.022 respectively).

We then examined tracts categorised into association/commissural fibres (gAF), thalamic radiations (gTR) and projection fibres (gPF). We found effects of probable MDD on measures of FA in two of the three groups of tracts. Probable MDD at principal and recurrent definitions showed smaller values in gAF (Probable MDD: β=-0.184, p_corrected_=0.010; Recurrent MDD: β=-0.170, p_corrected_=0.045) and gTR (Probable MDD: β=-0.159, p_corrected_=0.020; Recurrent MDD: β=-0.141, p_corrected_=0.068). No effect was found for gPF (Probable MDD: β=-0.115, p_corrected_=0.073; Recurrent MDD: β=-0.057, p_corrected_=0.401). The above findings were checked in self-declare depression, and the results were found to be similar (see supplementary materials, MDD definitions).

We then proceeded to compare FA values in the individual tracts between cases and controls. Initially, we tested the tracts controlling for hemisphere effects. Then we tested the interaction of hemisphere and probable MDD definitions on bilateral tracts to identify any lateralised effects. There was a significant interaction of hemisphere in superior longitudinal fasciculus for recurrent definition of probable MDD (β=0.117, p_corrected_=0.026). The left and right superior longitudinal fasciculi were therefore tested separately.

We found reduced FA in the left superior longitudinal fasciculus for both definitions of MDD versus controls (Probable MDD: β=-0.194, p_corrected_=0.025; Recurrent MDD: β=-0.221, p_corrected_=0.025) (Fig. 2, Table 2). No significant association was found with right superior longitudinal fasciculus (Principal MDD: Probable MDD: β=-0.057, p_corrected_=0.379; Recurrent MDD: β=-0.029, p_corrected_=0.684). Significant FA decrease was found in superior thalamic radiation and forceps major, but only for principal MDD definition (Probable MDD: β=-0.224, p_corrected_=0.009; β=-0.193 p_corrected_=0.025. Recurrent MDD: β=-0.179, p_corrected_=0.080; β=-0.133, p_corrected_=0.150 respectively for the two tracts). In order to check whether the decreased FA in the above tracts was due to global changes in gFA, the effect of MDD definitions was tested again with gFA included as a covariate (Table S6). Left superior longitudinal fasciculus remained significant in both definitions (Probable MDD: β=-0.194, p_corrected_=0.038; Recurrent MDD: β=-0.221, p_corrected_=0.025). Forceps major showed decreased FA in probable MDD definition (β=-0.193, p_corrected_=0.038) but not in recurrent MDD ( β=-0.133, p_corrected_=0.350). The effect MDD definitions on superior thalamic radiation didn’t reach significance after correcting for gFA (Probable MDD: β=-0.110, p_corrected_=0.162; Recurrent MDD: β=-0.077, p_corrected_=0.568). The above results of individual tracts turned null if outliers weren’t excluded, but the standard effect sizes were in similar trend (Table S7).

**Figure 2.**
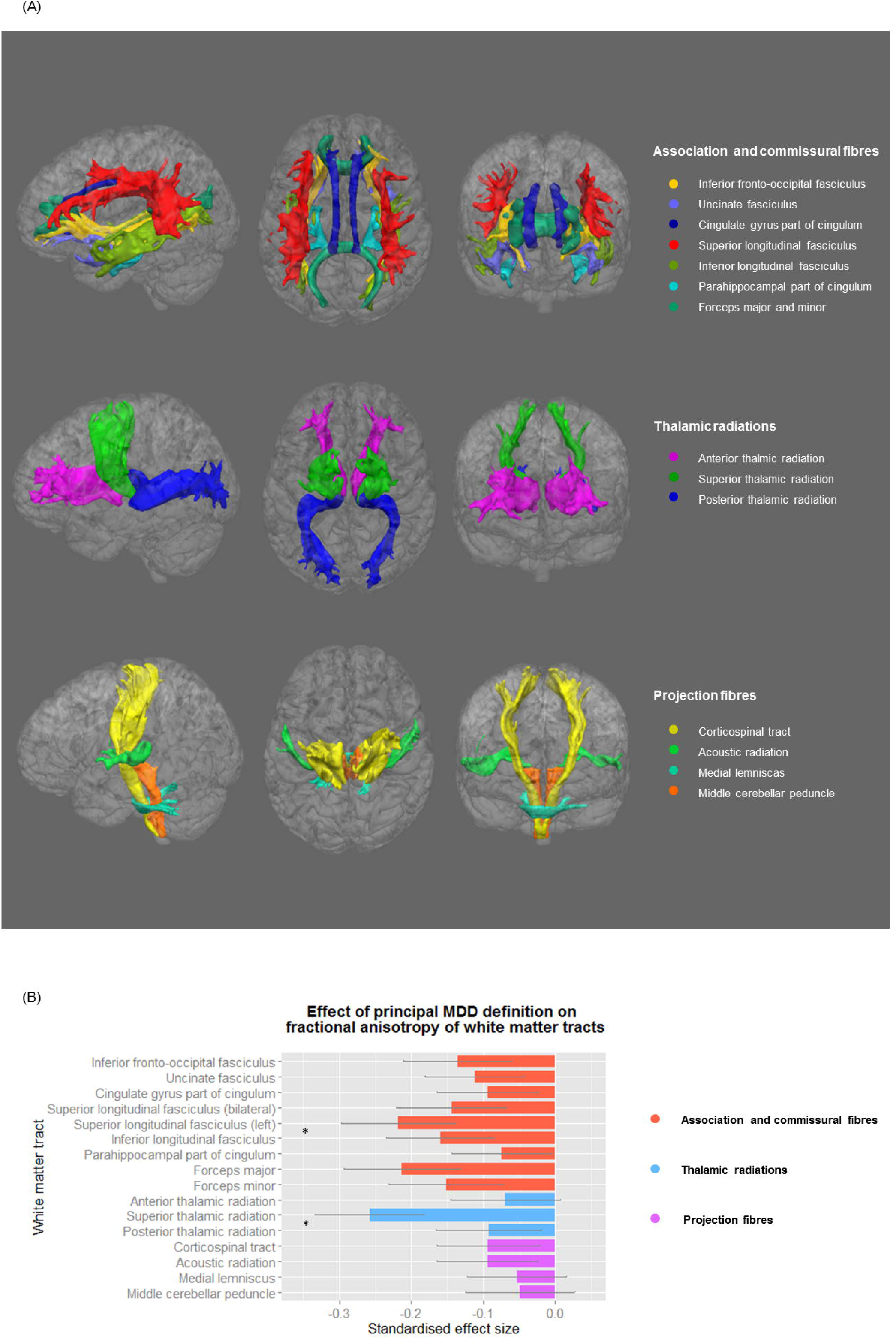
(A) White matter tracts in each anatomical subset in left, posterior and anterior view. (B) The effect of principal definition of probable MDD on FA value of tracts. Linear models were conducted, controlling the effect of age, age^2^, sex and assessment centre (and hemisphere for the tracts that have bilateral values). Left superior longitudinal fasciculus was presented because there was a significant interaction between recurrent MDD definition and hemisphere. Follow-up analysis showed a lateral effect of probable MDD definition on left superior longitudinal fasciculus. The x-axis shows the standardised effect size of MDD definition, and y-axis is the layout of the white matter tracts. The error bar represents standard deviation of mean.

**Table 2.**
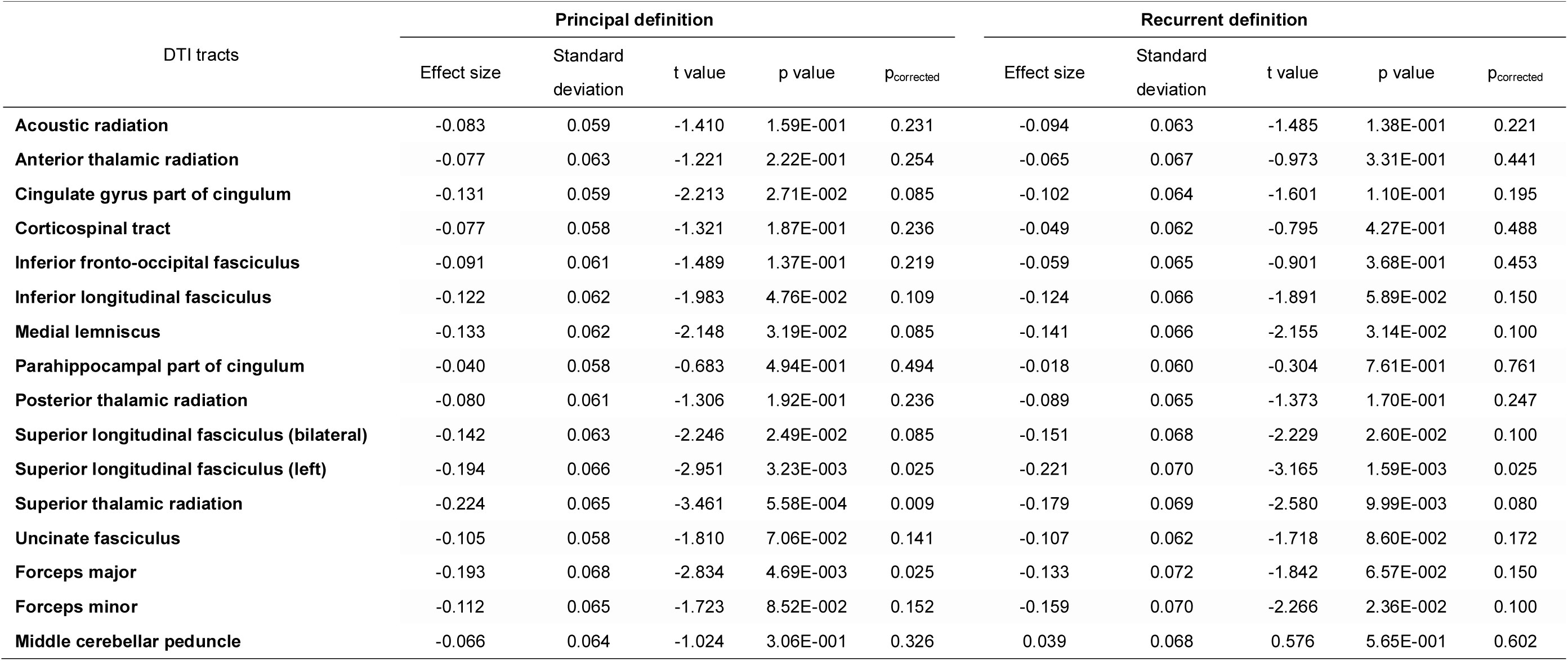
The effect of MDD definition on FA values of DTI tracts.

### Discussion

In the current study, we sought to determine whether MDD was associated with differences in subcortical grey matter volume or white matter integrity in a large imaging dataset from a single scanner of more than 8000 people, and among them over 1000 were included as cases and controls in the analyses for the present study. The sample sizes of MDD cases and controls included in the analysis of white matter integrity is by far the largest to our knowledge. Also, the present study considered two important brain structural modalities in two highly overlapping samples. Whilst we did not find any statistically significant subcortical volumetric differences between unaffected participants and individuals with probable MDD (using any of the definitions with increasing severity), we did find substantial evidence of reduced white matter integrity in MDD. This was seen globally, in two of the three categories of tracts (association/commissural fibres and thalamic radiation tracts), and in individual tracts (bilateral superior thalamic radiation, forceps major and left superior longitudinal fasciculus). Similar patterns of findings were seen for both principal and recurrent definition of depression with generally greater effect sizes in recurrent cases, with the exception of the localised differences in the superior thalamic radiation and forceps major.

Our study notably did not find evidence for bilateral hippocampal volume reduction as previously reported in the large collaborative meta-analysis of MDD^19^. We also did not find evidence of reductions in hippocampal volume when looking at recurrent MDD as published in the same study. The lack of subcortical volumetric differences associated with probable MDD diagnoses in the current study therefore does not support the widely held belief that there are subcortical volumetric changes associated with the disorder. There are several potential explanations for this. Firstly, the UK Biobank dataset included only community-dwelling, ambulant individuals who could independently complete the health and cognitive assessments, and attend the follow-up imaging assessments. This approach arguably selected MDD groups that were more well/better functioning but equally more representative of the general population than purely clinically ascertained samples. We also used a composite ‘probable’ MDD diagnosis that was based on self-report symptoms and hospital admission statistics, and the cases were selected based on self-report lifetime experience of probable depression. In contrast, many other studies previously used a structured clinical interview schedule, such as the Structured Clinical Interview for DSM-IV (SCID), to define MDD according to standard criteria. Some studies have specifically studied people who were certainly experiencing depression at the time of imaging assessment^33^. Whilst the probable MDD definitions used in the current paper were not based on an interview, they showed many of the same epidemiological and risk-factor associations as clinically defined cases^22,34^.

Although we do not report subcortical volume differences, we did find substantive evidence for robust deficits in both global and local white matter integrity. We found that MDD patients had global loss of FA which was also found to be reduced in association and commissural fibres as well as in thalamic radiations, but not in projection fibres. FA in these structures was also more severely reduced in the recurrent MDD patients. The above results indeed reflect previous findings from previous small-sample and meta-analytic studies^17,35,36^, while extending them to a more generalizable population-based cohort excluding potential methodological confounds as associated with the previous studies. A previous meta-analytic study that compared 231 MDD patients with 261 healthy participants found reduced FA in inferior longitudinal fasciculus, inferior fronto-occipital fasciculus, posterior thalamic radiation and corpus callosum, which belong to the association/commissural fibres and thalamic radiations^17^. Following the above study, another two recent meta-analyses found integrity reductions in the same categories, i.e. dorsal lateral PFC area, commissural fibres^35,37^. The global loss of FA in these regions could be the result of general neurodevelopmental alterations in MDD patients^38^, and findings within defined subsets of white matter tracts could reflect the neurological basis of MDD as a disconnection within an integrated network of cortex-cortex and cortical-limbic pathways^39^. The general FA reductions in groups of tracts is also consistent with findings from resting-state fMRI studies, which reported abnormalities in MDD populations in regional networks rather than just individual regions or structures^6,40^. The networks that derive from prefrontal cortex and thalamus has been found largely contribute to emotional and social cognition processes^38^. The reduced integrity in these groups of tracts may therefore reflect the repeatedly found impairment of emotion regulation^41,42^, reward processing^43^ and executive control^44^ in MDD populations.

In the tests of single white matter tracts, we found significantly altered integrity in left superior longitudinal fasciculus and superior thalamic radiation both in the overall MDD population and recurrent MDD patients. Reduction of left superior longitudinal fasciculus was notably larger in recurrent MDD patients. Reduction of integrity in forceps major was also found in MDD compared with healthy subjects, however showed no specific change of FA in recurrent MDD.

Superior longitudinal fasciculus, as a part of association fibres, connects prefrontal cortex and other lobes^45^. Small-sample studies have specifically reported reduced integrity in superior longitudinal fasciculus in various depressive samples, including elderly patients with depression^38,46^ depressive adolescents^47^ and adolescents with familial risk for depression^45^, compared with controls. Meta-analytic studies^35,48^ and a review^36^ also ascertained that the reduction of white matter integrity specifically in superior longitudinal fasciculus may be an important biomarker of the presence of depression. A recent study combined genetic and neuroimaging techniques found that people with higher polygenic risk of depression have greater loss of FA in superior longitudinal fasciculus^49^, suggesting that it may also therefore be a useful trait-related marker of risk. Loss of integrity in superior longitudinal fasciculus has also previously been reported to be associated with various cognitive dysfunctions, like working memory^50^ and attention^48^. Severity of depressive symptoms was also found correlate with FA loss in superior longitudinal fasciculus^51^. There is increasingly convincing evidence therefore that reduced integrity in superior longitudinal fasciculus might be an important feature of the neurobiology of MDD and may underlie impaired emotional process and cognitive abilities in MDD population^18^.

Another strength of the present study is that cross-modality assessment was conducted on both subcortical volumes and white matter integrity. Though the findings were largely found in white matter integrity instead of subcortical volumes, this is consistent with another cross-modality study by Sexton et al.(2012), which presented that no significant group difference was found between late-life depression and healthy control, whereas white matter integrity was reduced in many regions^16^. Another study on 358 people similarly found that depressive symptoms of elderly subjects also showed significant deficit in white matter, but not in grey matter measures^52^. The age range for the present study is from 40 to 70, which covers a notable range of elderly participants. This feature of our sample could be the reason why it showed similar contrast of findings between white matter and grey matter measurements.

Potential limitations of the current study should be considered, these include the absence of a face-to-face structured diagnostic interview schedule and the lack of hospital-based sampling. The large sample size may, however, overcome some of these difficulties and community based population sampling may yield more generalizable findings than those based on clinically ascertained samples alone^8,53^. The current investigation, by avoiding the combination of clinically and methodologically diverse samples, may also have ameliorated several important confounds such as differences due to different healthcare systems and illness related conditions including age of onset and duration of illness. Another factor of interest for future studies is the effect of hospital treatment. As studies have reported changes of depressive symptoms caused by medication or cognitive treatment^3^, investigates on the neurological effect of treatment should be conducted. The prevalence of the present study is lower than 10%, which is less than the prevalence of ~20% in overall sample of the cohort in the study by Smith et al. (2013)^22^. This was mainly due to the difference of sizes between the two samples. There were ~5500 participants in the sample with T1-weighted/DTI data, whereas over 30 times of people were included in the full cohort (N=172,751). This difference therefore supports the necessity of studying MDD in a large sample to minimise the bias of selecting study sample. A further potential limitation is that for the volumetric analysis we only focused on the subcortical volumes in the current study. We can therefore not exclude the possibility of cortical differences in MDD, including regional volume differences, as well as measures of cortical thickness and gyrification for example.

Our study presents a comprehensive comparison of brain structural changes related to MDD using the largest single sample available to date from a single scanner with uniform methodologies for clinical categorisation and scanning. We mainly report reductions of white matter FA in general latent measures of association and commissural fibres as well as thalamic radiations, and in left superior longitudinal fasciculus both in MDD and recurrent MDD. Future work would be potentially focusing on structural changes in cortical areas as well as richer stratification of MDD into informative biologically-based subgroups.

## Acknowledgements

This study is supported by a Wellcome Trust Strategic Award “Stratifying Resilience and Depression Longitudinally” (STRADL) (Reference 104036/Z/14/Z).

We thank the UK Biobank participants for their participation, and the UK Biobank team for their work in collecting and providing these data for analysis. This research was conducted, using the UK Biobank Resource under approved project 10279, in The University of Edinburgh Centre for Cognitive Ageing and Cognitive Epidemiology (CCACE) (http://www.ccace.ed.ac.uk), part of the cross-council Lifelong Health and Wellbeing Initiative (MR/K026992/1). XS receives support from China Scholarship Council. HCW is supported by a JMAS SIM fellowship from the Royal College of Physicians of Edinburgh and by an ESAT College Fellowship from the University of Edinburgh. SRC was supported by Medical Research Council grant MR/M013111/1. IJD and DCL are supported by the Medical Research Council award to CCACE (MR/K026992/1). IJD is additionally supported by the Dementias Platform UK (MR/L015382/1), and he and SRC by the Age UK-funded Disconnected Mind project (http://www.disconnectedmind.ed.ac.uk). DJS is supported by an Independent Investigator Award from the Brain and Behaviour Research Foundation (21930). Part of the work was undertaken in The University of Edinburgh Centre for Cognitive Ageing and Cognitive Epidemiology (CCACE), funding from the Biotechnology and Biological Sciences Research Council (BBSRC) and Medical Research Council (MRC) is gratefully acknowledged. Age UK (The Disconnected Mind project) also provided support for the work undertaken at CCACE.

## Author contributions

AMM, HCW, SRC, LMR and XS contributed to the design of the study, analysis of the data, and writing the manuscript. MJA was involved in curating the data. IJD, DCL, DJS and MEB were involved in the conception of the study and overseeing analysis methodology. UK Biobank collected all data and was involved in the preprocessing of imaging data. All authors discussed and commented on the manuscript.

